# Diagnostic value of blood gene expression-based classifiers as exemplified for acute myeloid leukemia

**DOI:** 10.1101/382143

**Authors:** Stefanie Warnat-Herresthal, Konstantinos Perrakis, Bernd Taschler, Matthias Becker, Lea Seep, Kevin Baßler, Patrick Günther, Jonas Schulte-Schrepping, Kathrin Klee, Thomas Ulas, Torsten Haferlach, Sach Mukherjee, Joachim L. Schultze

## Abstract

**ABSTRACT:** Acute Myeloid Leukemia (AML) is a severe, mostly fatal hematopoietic malignancy. Despite nearly two decades of promising results using gene expression profiling, international recommendations for diagnosis and differential diagnosis of AML remain based on classical approaches including assessment of morphology, immunophenotyping, cytochemistry, and cytogenetics. Concerns about the translation of whole transcriptome profiling include the robustness of derived predictors when taking into account factors such as study- and site-specific effects and whether achievable levels of accuracy are sufficient for practical use. In the present study, we sought to shed light on these issues via a large-scale analysis using machine learning methods applied to a total of 12,029 samples from 105 different studies. Taking advantage of the breadth of data and the now much improved understanding of high-dimensional modeling, we show that AML can be predicted with high accuracy. High-dimensional approaches - in which multivariate signatures are learned directly from genome-wide data with no prior biological knowledge - are highly effective and robust. We explore also the relationship between predictive signatures, differential expression and known AML-related genes. Taken together, our results support the notion that transcriptome assessment could be used as part of an integrated genomic approach in cancer diagnosis and treatment to be implemented early on for diagnosis and differential diagnosis of AML.

**One Sentence Summary:** Blood gene expression data and machine learning were used to develop robust and accurate classifiers for diagnosis and differential diagnosis of acute myeloid leukemia based on analysis of more than 12,000 samples derived from more than 100 individual studies

## Introduction

Recommendations for diagnosis and management of malignant diseases are organized by international expert panels. For example, the first edition of the European LeukemiaNet (ELN) recommendations for diagnosis and management of acute myeloid leukemia (AML) in adults were published in 2010 *(1)* and recently revised in 2017 *(2)*. Based on the most recent DNA-sequencing results, such as those derived from The Cancer Genome Atlas (TCGA), AML can be subdivided into multiple subclasses *(3–11)*. While subclassification is already based on genomic information, according to the ELN recommendations primary and differential diagnosis still relies on classical approaches including assessment of morphology, immunophenotyping, cytochemistry and cytogenetics *(2)*.

This is surprising considering that in a pioneering study almost two decades ago Golub et al. demonstrated that AML could be identified by combining gene expression profiling (GEP) and computational class prediction *(12)*. Admittedly, this initial study was very small and without a true validation cohort, but subsequent work promised to fundamentally change primary and differential diagnosis of AML by assessing the transcriptome *(13–17)*. Furthermore, it had been suggested that GEP could be utilized to define leukemia subtypes and derive useful predictive gene signatures *(18, 19)*. A decade later, the International Microarray Innovations in Leukemia Study Group even proposed GEP by microarray analysis to be a robust technology for the diagnosis of hematologic malignancies with high accuracy *(20)*. The utility of GEP by RNA-sequencing has been also demonstrated for other tumor entities for example breast cancer *(21–23)*, bladder or lung cancer *(24, 25)*. Also, in AML research large RNA-seq datasets have been described in the meantime *(26–30)*. However, despite these promising results and data, GEP did not translate into clinical practice for AML diagnostics.

In parallel, a series of advances in machine learning (ML) and computational statistics have transformed our understanding of prediction using high-dimensional data. A variety of approaches are now an established part of the toolkit and for some models (including sparse linear and generalized linear models), there is a rich mathematical theory concerning their performance in the high-dimensional setting *(31)*. In a nutshell, the body of empirical and theoretical research has shown that learning predictive models over large numbers of variables is often feasible and remarkably effective. In applied machine learning, there has been a deepening understanding of practical issues, e.g. relating to the transferability of predictions across contexts *(32)*, that is very relevant to the clinical setting.

Based on these developments in the data sciences and the increasing availability of GEP data derived from peripheral blood including AML, we sought to re-assess the potential for translation of transcriptomic data into AML diagnostics. To this end, we built the probably largest reference blood GEP dataset comprising 105 individual studies with in total more than 12,000 patient samples. We applied high-dimensional machine learning approaches to build genome-wide predictors in an unbiased, entirely data-driven manner, and tested predictive accuracy in held-out data. We carried out extensive tests designed to address specific concerns relevant to practical use, including the case of transferring predictive models between entirely disjoint studies (that could be subject to batch effects or other unwanted variation) and even between transcriptomic platforms. Our results show that combining machine learning and blood transcriptomics can yield highly effective and robust classifiers. This supports the notion that GEP should be considered for AML diagnostics and tested directly against currently recommended classical approaches.

## Results

### Establishment of a unique GEP dataset for classifier development

We hypothesized that the determination and comprehensive evaluation of GEP-based AML classifiers requires large datasets, should include samples from many sources to mimic the situation in real-world deployment and should include several technical platforms to better understand their influence on classifier performance. To achieve these goals, we wanted to include the largest number of peripheral blood mononuclear cells (PBMC) or bone marrow samples possible and therefore systematically searched the National Center for Biotechnology Gene Expression Omnibus (GEO) database for PBMC and bone marrow studies (Fig. 1A). We identified 153,922 datasets, of which 111,632 also contained human samples. To include only whole sample series and to avoid duplicate samples, we filtered for GEO series (GSE) and excluded so-called super series, which resulted in 6800 studies. Of those, 5320 contained bone marrow and PBMC samples, of which 2715 were human studies. We then focused the analysis on studies with samples analyzed on one of three platforms including the HG-U133A microarray, the HG-U133 2.0 microarray and Illumina RNA-seq. Next, duplicated samples and studies working with pre-filtered cell subsets were excluded. This study search strategy resulted in 105 studies with a total of 12,029 samples (Fig 1) including 2,500 samples assessed by HG-U133A microarray (Dataset 1), 8,348 samples by HG-U133 2.0 microarray (Dataset 2) and 1,181 samples by RNA-seq (Dataset 3). In total, the dataset contained 4,145 AML samples of diverse disease subtypes and 7,884 other samples derived from healthy controls (*n*=904), patients with acute lymphocytic leukemia (ALL, *n*=3,466), chronic myeloid leukemia (CML, *n*=162), chronic lymphocytic leukemia (CLL, *n*=770), myelodysplastic syndrome (MDS, *n*=267) and other non-leukemic diseases (*n*=2,312) (Fig. 1B, Fig S1). Unless otherwise noted, all samples derived from AML patients are referred to as cases and non-AML samples as controls. We additionally considered differential diagnosis, in which case the controls comprised non-AML leukemias. According to the three platform types, the whole sample cohort was divided into three datasets referred to as datasets 1, 2 and 3 (Table S1, Fig. S1).

**Figure 1:**
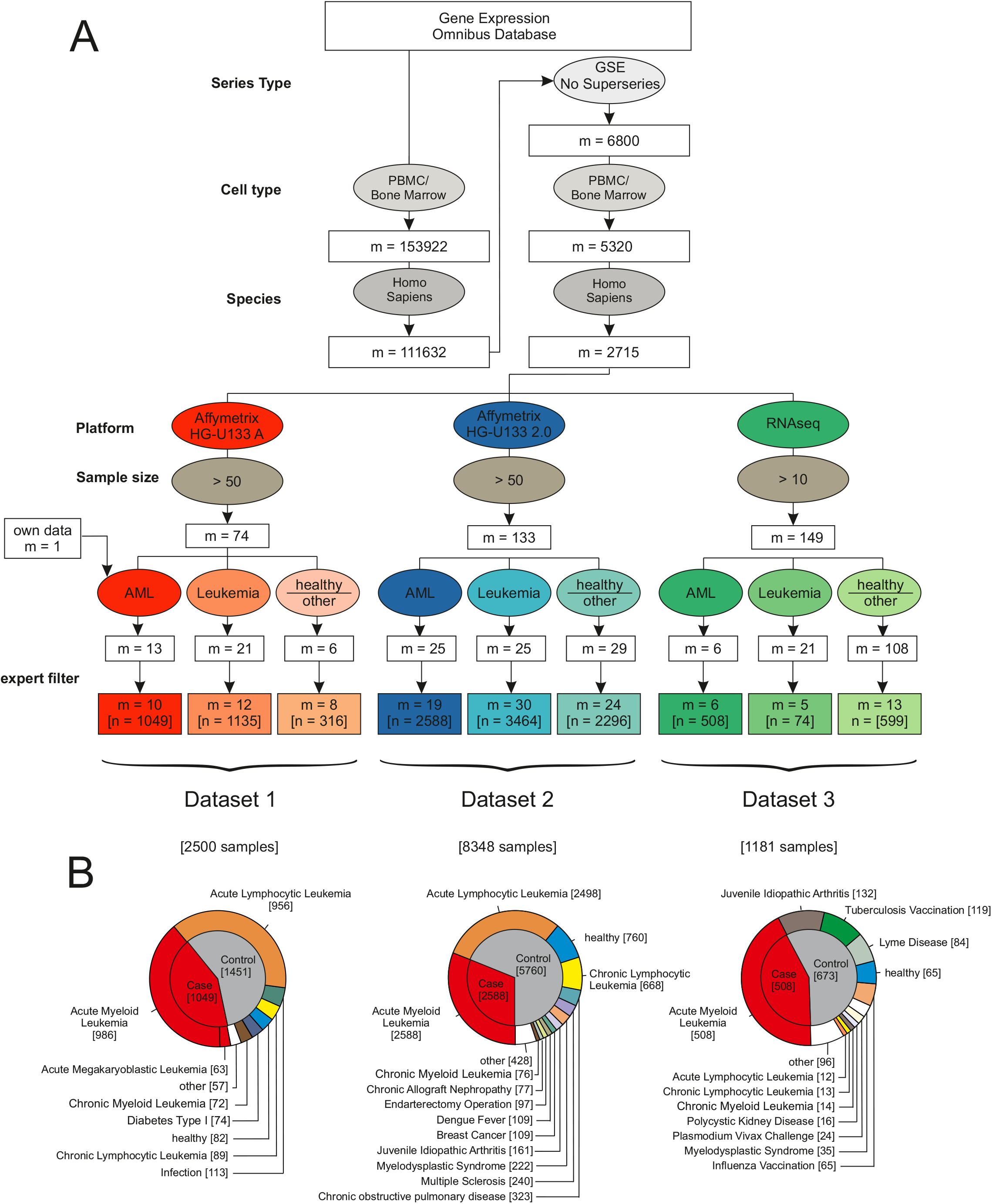
Establishing datasets for the largest AML meta-study to date. (**A**) Flowchart for the inclusion of studies. The gene expression omnibus (GEO) database was systematically searched for GEO Series of human PBMC and bone marrow samples processed with microarray platforms (Affymetrix HG-U133A and HG-U133 2.0) or next generation RNA sequencing (RNA-seq) data. These data were filtered for inclusion of either AML samples, samples of other leukemia and inclusion of healthy samples or other diseases. After manual revision and exclusion of duplicates and experiments using sorted cell populations (“expert filter”), the data was combined and normalized independently for each dataset. In total, the datasets contained 105 studies and 12,029 samples. (**B**) Detailed overview of the three datasets established in this study after filtering as given in (A). Dataset 1 (Affymetrix HG-U 133 A) contained 986 AML and 63 AMKL samples (cases; a total of 1049 samples) and 1451 non-AML samples with the majority (956) being ALL samples. Dataset 2, being the largest dataset in this study (8348 samples), consisted of 2588 AML and 5760 non-AML samples of leukemic and non-leukemic diseases, respectively. Dataset 3 (RNA-seq) comprised 508 AML samples with 673 control samples of mostly non-leukemic diseases.

### Effective AML classification using high-dimensional models

Here, we sought to assess classification of AML vs non-AML. Microarray data were RMA normalized while RNA-seq data was normalized as implemented in the R package DESeq2 *(33)*. For further analysis and better comparison between the different datasets, we trimmed the data to 12,708 genes identified within all datasets (Fig. 2A). The size of the test set was 20% of the total sample size and random sampling of training and test sets was repeated 100 times. As main performance metrics, we considered (held-out) accuracy, sensitivity and specificity. Classification was performed using *l*_1_-regularized logistic regression (the lasso; see also below).

**Figure 2:**
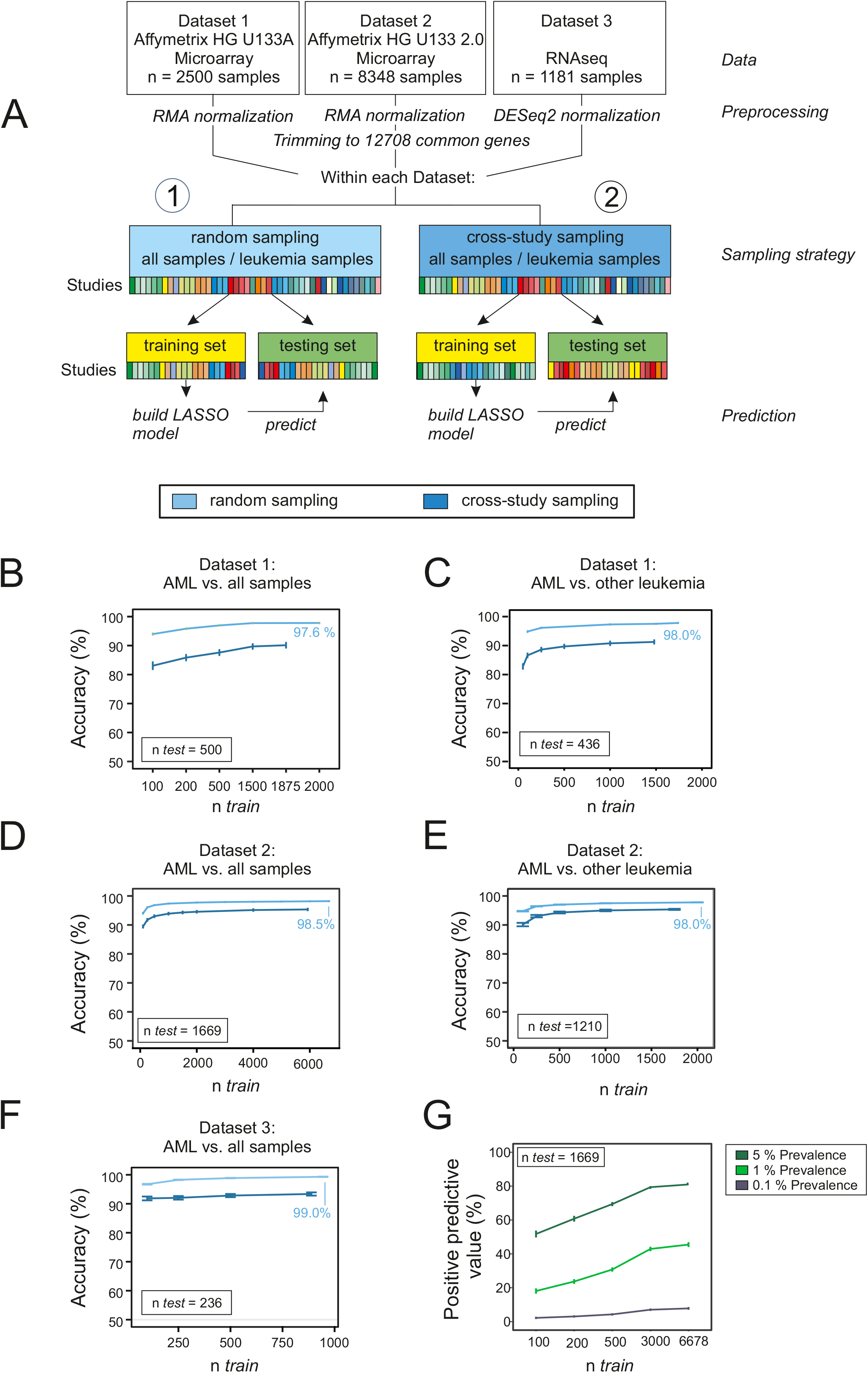
Prediction of AML in random and cross-study sampling scenarios. (**A**) Workflow showing preprocessing and sampling steps for each dataset and two sampling strategies (random and cross-study sampling) for splitting the data into training and test sets. Classification accuracies based on the lasso model of AML versus all other samples and for both sampling strategies are shown for dataset 1 (**B**), dataset 2 (**D**) and dataset 3 (**F**). Classification accuracies for the differential diagnosis case (AML versus other leukemic samples, namely AML, ALL, CML, CLL and MDS) for both sampling strategies are shown for dataset 1 (**C**) and dataset 2 (**E**). Mean accuracies of the lasso models are shown as a function of the training sample size *n_train_*. Results are over 100 random training and test sets, with error bars indicating the standard deviation. (**G**) Positive predictive value (PPV) as a function of *n_train_*, corresponding to the setting (**D**) and assumed prevalence of 0.1%, 1% or 5% (see text).

First, we included all non-AML samples, including healthy controls and non-leukemic diseases, among the controls (Fig. 2B, D, F, light blue lines). The goal was to classify unseen samples as AML or control. To understand how much data is needed in this setting, we plotted learning curves showing the test set accuracy as a function of training sample size *n_train_*. For each gene expression platform, this was done by randomly subsampling *n_train_* samples and testing on held-out test data with fixed sample size *n_test_* (as shown). We see that prediction in this setting is already highly effective with a small number of training samples, although accuracy still increases with increasing *n_train_* (note that the total number of samples and hence range of *n_train_* differs by platform).

In many clinical settings, the control group does not contain healthy controls, but rather related diseases. To test effectiveness in a differential diagnosis setting, we repeated the experiments but with controls sampled only from other leukemic diseases, such as ALL, CLL, CML and MDS (Fig. 2C, E, Fig. S2–4). We observed similar prediction results, which indicated that prediction accuracy is not only due to large differences between AML and non-leukemic conditions.

For diagnostic utility, the positive predictive value (PPV; the probability of disease given a positive test result) is an important quantity. The PPV depends not only on sensitivity and specificity but also on prevalence, as it is harder to achieve a high PPV for a condition that is rare in the population of interest. Although we found high accuracy, sensitivity and specificity already at moderate *n_train_*, depending on the use case, this could still imply that large training sample sizes would be useful to reach higher PPVs. For example, the predictive gains in increasing *n_train_* from the lowest to highest values indicated in Fig 2D, which is for the dataset with largest total sample size, correspond to a *doubling* of PPV from ~20% to ~40% at an assumed prevalence of 1% (Fig 2G). This illustrates the fact that although after a certain point increasing *n_train_* tends to increase accuracy only slowly, the gains, even if small in absolute terms, can be highly relevant with respect to PPV in low-prevalence settings.

In additional experiments we considered performance of nine different classification methods (Figs S2–4). We could predict AML with high accuracy with all tested classification algorithms on microarray platforms (Fig. S2 & S3). For RNA-seq data, the lasso and random forests were able to predict with high sensitivity and specificity (Fig. S4). Lasso-type methods have several advantages, including extensive theoretical support and interpretability, so we focused on these as our main predictive tool. Deep neural networks provided comparable prediction performance to the lasso (data not shown), but we preferred the latter in this setting again due to interpretability, since the lasso provides explicit variable selection, facilitating model interpretation.

### Assessing the effect of cross-study-variation on predictive performance

Microarray data and data generated by high-throughput sequencing are both known to be susceptible to batch effects *(34)*. More generally, diverse study-specific effects and sources of study-to-study variation can pose problems in the context of predictive tests for clinical applications. Predictors that perform well within one study may perform worse when applied to data from new studies *(35)* with implications for practical generalizability.

The results above spanned data from multiple heterogeneous studies. Provided training and test data are sampled in the same way, such heterogeneity does not necessarily pose problems for classification, as evidenced above. However, if the training and test data are from entirely different sites/studies (rather than randomly sampled from a shared pool), then the impact of batch/study effects may be more serious. We took advantage of the large number of studies in our dataset to sample training and testing sets in such a way that they were mutually disjoint with respect to studies. That is, any study that was at all represented in the training set was entirely absent from the test set and *vice versa* and we use the term *cross-study* to refer to this strictly disjoint case. Results are shown in Fig 2B,D,F (dark blue lines). As expected, performance was worse in the cross-study setting than under entirely random sampling (light blue lines). However, in the dataset with the largest sample size (dataset 2, platform HG-U133 2.0; Fig. 2D) we see that the performance in the cross-study case gradually catches up to the random sampling case with only a small gap at the largest *n_train_*. The other two datasets have smaller total sample sizes so never reach comparable training sample sizes. Note that we did not carry out any batch effect removal using tools such as SVA *(36)* or RUV *(37)* and in that sense our results are conservative. Despite the availability of these and other tools for batch effect correction, it is difficult to be fully assured of the removal of unwanted variation in practice. Our intention here was not to remove between-study variation but rather to (conservatively) quantify its effects on accuracy.

Due to the large number of studies included in our analysis, we were able to carry out an entirely disjoint cross-study analysis also for the differential diagnosis case. These results are shown in Fig 2C,E (dark blue lines; cross-study sampling for differential diagnosis was not possible using dataset 3 due to lack of samples, see Fig. S1) and are broadly similar, also across different classification algorithms (S5-S7).

### Translation of classifiers across technical platforms

Over the long term, clinical pipelines must cope with changes in technological platforms. It is therefore relevant to understand to what extent predictors can generalize not just between studies, but between different platforms. In other words: is it possible to take a model learned on data from platform *A* and deploy it using unseen data from platform *B*? To address this question, we constructed AML vs non-AML training and test sets in a *cross-platform* manner, i.e. training on one platform and testing on another (Fig. 3A). That is, a model was learned using data from one platform and then this model – used “as is”, with no further fine tuning – was used to make predictions using expression data from a different platform. We see that classification accuracy varies greatly. Classifiers that were trained on HG-U133 A (dataset 1) work well when tested using data generated with the more advanced microarray HG-U133 2.0 (dataset 2) (Fig. 3B) and models trained on HG-U133 2.0 data can predict well using RNA-seq data (dataset 3) (Fig. 3D). However, models trained naively on HG-U133 A data cannot predict using RNA-seq data (Fig. 3F).

**Figure 3:**
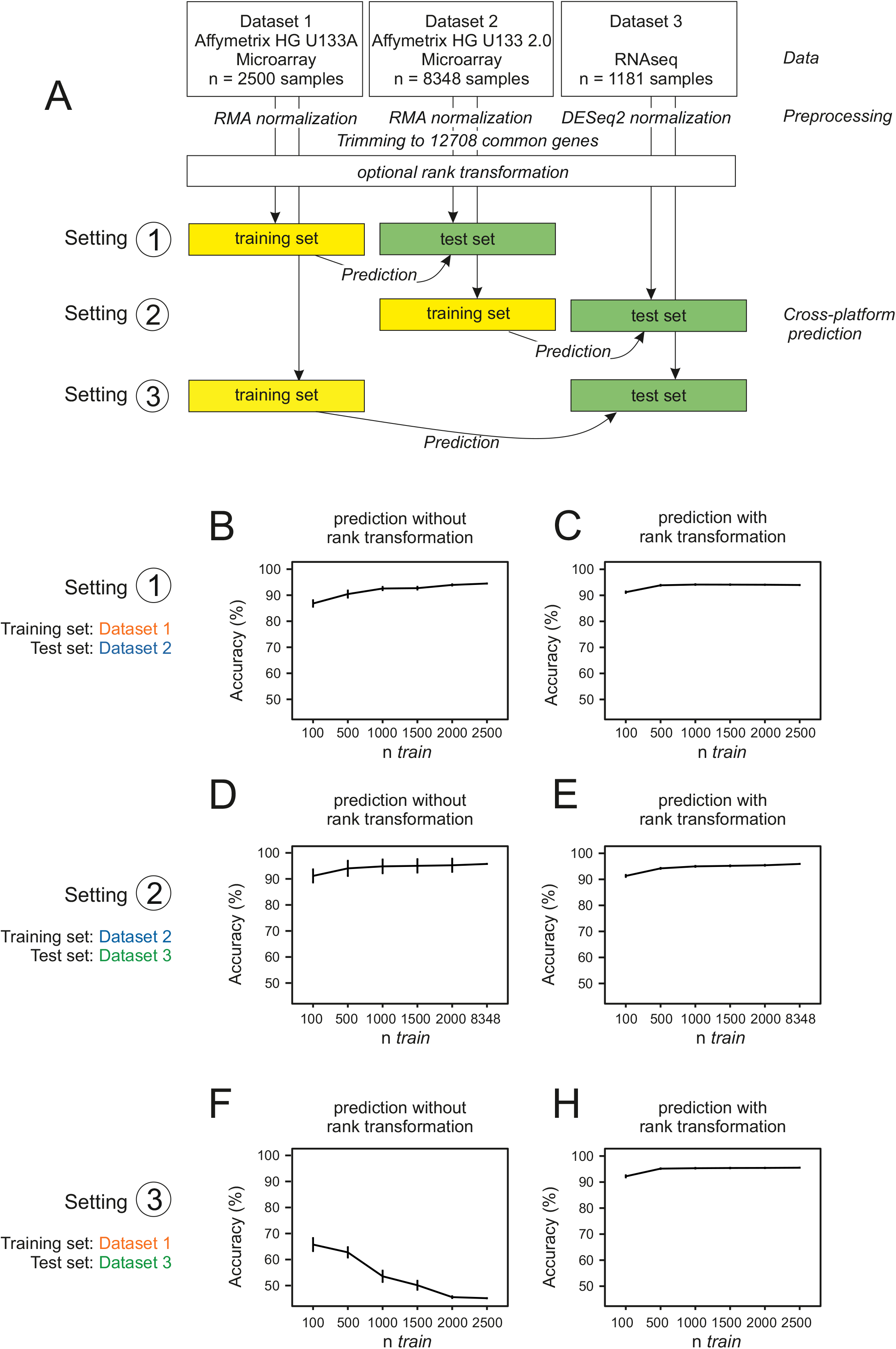
Translating predictive signatures across technological platforms. (**A**) Datasets were normalized individually and trimmed to 12708 common genes. The classifiers were trained on subsamples of different sizes on one platform and tested on all samples of another platform. Classification accuracies are shown as a function of training sample size (*n_train_*) with rank transformation (**C**, **E**, **H**) and without rank transformation (**B**, **D**, **F**). For the former case, the training and test datasets (from different platforms) were separately rank transformed (see text for details).

To explore the utility of simple transformations in this context, we then performed a rank transformation to normality on all datasets. This is among the simplest and best known data transformations and has previously been shown to increase the performance of prognostic gene expression signatures and can even outperform more complex variance-stabilizing approaches *(38)*. With this approach, we reached very good overall performance across all platforms under study (Fig 3 C, E, H). Note that the transformation is simply applied to each dataset independently and could be easily deployed in any practical use-case without any need for prior input into e.g. cross-platform designs such as inclusion of control samples.

### Predictive signatures and AML biology

The predictive models derived from the lasso and used above are sparse in the sense that they automatically select a small number of genes to drive the prediction. The genes are selected in a unified global analysis, rather than by differential expression on a gene-by-gene basis. From a statistical point of view, global sparsity patterns for prediction and gene-by-gene differential expression are different criteria. Furthermore, a good set of genes for prediction need not be mechanistic (in the sense of constituting causal drivers of the disease state). We therefore sought to understand the relationship between differential expression (DE), known mechanisms and predictive gene signatures.

Using dataset 2 (the largest dataset) we compared DE and the sparse predictive models. We performed DE analysis using the whole dataset and compared the results with the set of genes in the lasso model (“lasso genes”) based on the same data (Fig 4A). 506 genes were differentially expressed (“DE genes”), of which 26 were associated with the disease ontology term or KEGG pathway for AML (“AML-related genes”). Of the 141 Lasso genes, 7 genes were leukemia-related and 46 were DE genes, meaning that many of the lasso genes were not differentially expressed, as clearly seen when overlaying the lasso gene selection on a volcano plot (Fig 4A). This underlines the fact that differential expression and predictive value in a signature sense are different criteria.

**Figure 4:**
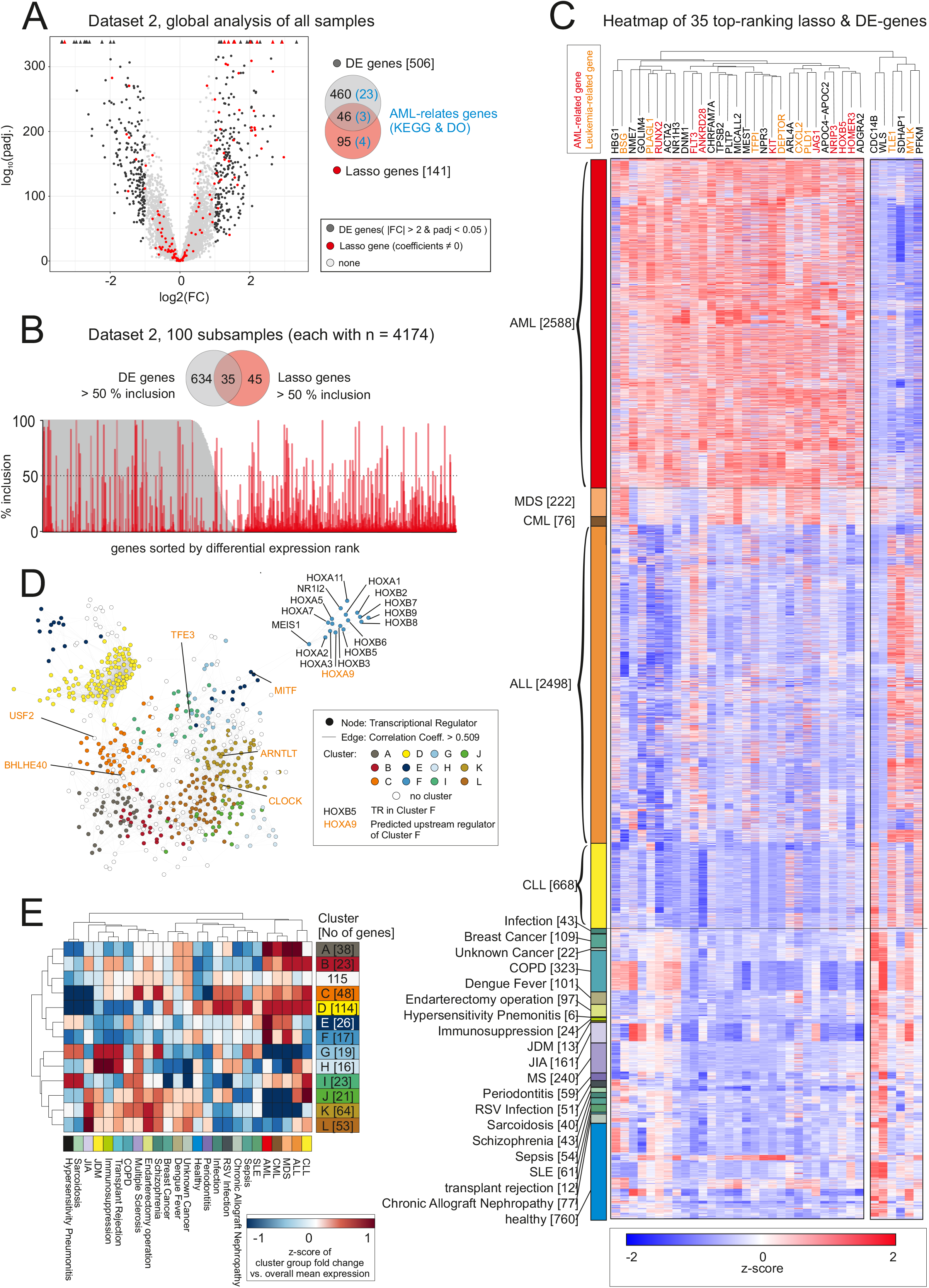
Predictive signatures and AML biology. (**A**) Volcano plot of global differentially expressed (DE) genes and genes of the lasso model (“lasso genes”) in dataset 2 and Venn diagram indicating the overlap of both gene sets and the genes included in the KEGG-pathway or the disease ontology term “AML”. (**B**) Inclusion plot of DE genes and lasso genes in 100 random permutations of dataset 2. The plot is sorted according to DE gene rank and a Venn diagram shows the overlap between genes with a minimum of 50 % inclusion. (**C**) Heatmap and hierarchical clustering of z-scaled expression values of 35 genes with > 50 *%* inclusion both in lasso and DE genes, as shown in (**B**). Genes with known associations with AML are marked red, genes associated with other types of leukemia are labeled in orange. (**D**) Co-expression network of 613 transcriptional regulators with 13 clusters as defined by 1000x infomap clustering. Genes of cluster F are labeled in black, predicted upstream regulators are shown in orange. (**E**) Heatmap and hierarchical clustering of the mean expression within clusters vs. the overall mean expression for all diseases.

Next, we extended this analysis to focus on DE and lasso genes whose selection was robust to data subsampling. This was done by subsampling half the dataset randomly 100 times and in each such subsample carrying out the full DE and lasso analyses. For the lasso this type of approach has been studied under the name stability selection *(39)*. DE and lasso genes were then scored according to the frequency with which they appeared among the 100 rounds of selection (Fig 4B). Thus, an inclusion score of 100% for a DE gene means that the gene is selected as differentially expressed in all 100 iterations and similarly for the lasso genes. In total, 669 genes passed the DE cutoffs in a least 50% of the iterations, while 80 genes were called in at least 50% of the iterations by the lasso-model (Fig. 4B). Of these genes, 35 were called according to both criteria. These results show that even among the genes that are included in the lasso models with high frequency (i.e. those genes that are robustly selected for prediction), many are not differentially expressed. To look at the relationship between differentially expressed genes and prediction from another perspective, we entirely *excluded* the 1000 most differentially expressed genes, and re-learned predictive signatures. Classification accuracy was essentially unaffected, demonstrating the robustness of high-dimensional predictive analyses to the absence of even the highest-ranked DE genes.

To better understand the biological relevance of these 35 genes for AML, we visualized the top ranked genes over all 8,348 samples within the dataset by hierarchical clustering of z-transformed expression values (Fig. 4C). We identified one distinct cluster of genes with the majority of genes being elevated in AML compared to other leukemias and non-leukemic samples (cluster 1, *n*=29). While we identified several well-known AML-related genes (gene name in red color) such as the KIT Proto-Oncogene Receptor Tyrosine Kinase (KIT) *(40–42)*, RUNX2 *(43)*, and FLT3 *(44, 45)* in this cluster, many genes have not yet been linked to AML biology and, although not the focus of the present paper, further study of these genes may be interesting from a mechanistic point of view. Within the other cluster (cluster 2, *n*=6 genes), genes had reduced expression values in AML compared to other leukemias and two of these genes have been linked to other types of leukemias (gene names in orange color).

Numerous transcriptional regulators (TRs) were among the top 35 ranked genes, including one HOX transcription factor. To further elucidate this finding, we performed a co-expression network analysis based on all TRs (Fig. 4D) and used infomap clustering to further explore the network (Fig. 4E). Three central clusters were defined that were further subdivided into smaller clusters. To find out whether clusters of co-expressed TRs were specific for distinct disease types, we visualized the fold change of the mean expression of every group versus the mean over all samples (group fold change) for each cluster and each disease in a heatmap (Fig. 4E), which indicated pan-leukemia clusters as well as cluster specific for AML and the closely related MDS (dark blue cluster, n = 26 genes and middle blue cluster, n = 17 genes). Interestingly, one of those AML-related clusters was particularly populated by HOX TFs (Fig. 4D) of which HOXA9 has been linked to poor AML prognosis *(46–48)*. To determine a potential master regulator, we performed TF motif enrichment analysis on this cluster and identified HOXA9 as one of the most prominent candidate master regulators of this AML-associated TF cluster.

## Discussion

Despite the pioneering studies by Golub and others (12–17) suggesting high potential value of GEP for AML diagnosis and differential diagnosis, the most current recommendations for this disease still center on classical approaches including assessment of morphology, immunophenotyping, cytochemistry, and cytogenetics *(2)*. By making use of more than 12,000 samples from more than 100 individual studies, we provided evidence that the combination of large GEP data with statistical and machine learning approaches allows for the development of robust disease classifiers. Such classifiers could be used for diagnosis and differential diagnosis of this deadly disease. Considering the increased utilization of whole genome and transcriptome sequencing in the management of cancer patients, we propose that clinical application of GEPand machine learning-based classifiers for diagnosis including differential diagnosis - instead of, or alongside, the more classical techniques - needs to be re-evaluated. This is in line with previous suggestions by the International Microarray Innovations in Leukemia Study Group *(20)*. We suggest that similar analyses may be useful for other diseases, when analyzing whole blood or PBMC-derived gene expression profiles.

We analyzed the largest collection of data so far in AML transcriptomics with the aim of understanding and addressing bottlenecks in the way of clinical deployment of gene expression-based machine learning tools for diagnosis. We considered a range of practical concerns, including differential diagnosis, cross-study issues and prediction across different technological platforms. We found that highly accurate prediction is possible in all scenarios considered. From a technical perspective, all the approaches we employed could be deployed at scale. We found also that in most cases good test results were achieved with relatively few training samples. The exception was for cross-study prediction, where a larger number of samples was needed to match the predictive performance seen in the (easier) random sampling case.

Site- and study-specific effects may be relevant for clinical applications. This is due to the fact that a classifier once learned might be deployed in a range of new settings (sites, regions) that could lead in a number of ways to unwanted variation. If training and test sets are very different, this can impact performance. Nevertheless, we saw that given sufficiently large training samples, cross-study prediction was effective. This suggests that while standardizing data collection as much as possible is of course desirable, given strong enough signals it may be possible to deliver useful predictions even in the face of unwanted and perhaps unknown variability. Nevertheless, in clinical applications of predictive models it will be important to continually track performance even after deployment.

All our models were learned in an unbiased manner, directly from the full transcriptome data with no prior biological knowledge or any pre-selection of genes. We showed that genes relevant for prediction were often not differentially expressed and that prediction was robust to removal of highly differentially expressed genes. These observations illustrate two points of relevance to clinical applications. First, that for prediction it can be more fruitful to consider signatures derived in data-driven, genome-wide fashion than to think in terms of single genes or differential expression. Second, that high-dimensional analyses, although complex relative to more classical methods, can be highly predictive as well as robust to the presence or absence of specific genes. Taken together, our results underline the immense value of making GEP data publicly available, allowing for new and large-scale multi-study analyses. Further, we believe that the application of statistical and machine learning approaches based on sequencing data to identify gene signatures for certain diseases such as AML will become part of recommendations for diagnosis and management of AML. We envision that combining whole genome and transcriptome analysis based on machine learning algorithms will ultimately allow diagnosis, differential diagnosis, subclassification and outcome prediction in an integrated fashion.

## Methods

### Review and selection of gene expression data

All data sets published in the National Center for Biotechnology Information Gene Expression Omnibus (GEO, Barrett et al., 2012) on 20 September 2017 were reviewed for inclusion in the present study. Basic criteria for inclusion were the cell type under study (human peripheral blood mononuclear cells (PMBCs) and/or bone marrow samples) as well as the species *(Homo sapiens)*. Furthermore, we excluded GEO SuperSeries to avoid duplicated samples (Table S1). We filtered the datasets for data generated with Affymetrix HG-U133 A microarrays, Affymetrix HG-U133 2.0 microarrays and high-throughput RNA sequencing (RNA-seq) and excluded studies with very small sample sizes (< 50 samples for microarray and < 10 samples for RNA-seq data). We then applied a disease-specific search, in which we filtered for acute myeloid leukemia, other leukemia and healthy or non-leukemia-related samples.

The results of this search strategy were then internally reviewed and data were excluded based on the following criteria: (i) exclusion of duplicated samples, (ii) exclusion of studies that sorted single cell types (e.g. T cells or B cells) prior to gene expression profiling, (iii) exclusion of studies with inaccessible data. Other than that, no studies were excluded from our analysis. In addition, we included one unpublished dataset (in dataset 1). The above steps gave rise to the data referred to above as **dataset 1** (Affymetrix HG-U133 A microarrays), **dataset 2** (Affymetrix HG-U133 2.0 microarrays) and **dataset 3** (RNA-seq).

### Pre-processing

All raw data files were downloaded from GEO. For normalization, we considered all platforms independently, meaning that normalization was performed separately for the samples in dataset 1, 2 and 3, respectively. Microarray data (datasets 1 and 2) were normalized using the robust multichip average (RMA) expression measures *(49)*, as implemented in the R package affy *(50)*. RNA-seq data (dataset 3) was normalized with the R package DeSeq2 using standard parameters *(33)*. In order to keep the datasets comparable, we filtered the data for genes annotated in all three datasets, which resulted in 12.708 genes. No filtering of low-expressed genes was performed. All scripts used in this study for pre-processing are provided as a docker container on GitHub *(51)*.

### Prediction

Prior to classification, data sets were split into non-overlapping training and test data. All classification tasks were performed in the programming language R *(52)*. All main results were obtained using *l*_1_-penalized logistic regression using the package glmnet *(53)*. Non-zero coefficients were extracted for feature ranking (Figure 4). The sparsity parameter was set using 10-fold cross-validation (using training set data only). To assess predictive performance, accuracy, sensitivity and specificity were calculated as well as positive predictive value (PPV) under several prevalence scenarios. For assessing the performance of support vector machines (SVMs), we used the R package e1071 for SVMs (linear, radial, polynomial and sigmoid kernels) *(54)*. The R package randomForest was used for random forest classification *(55)*. K nearest neighbors classification was done using the knn function implemented in the class package in R *(56)*. Linear discriminant analysis was performed with the lda function implemented in the R package MASS *(56)*. For RNA-seq data, features with zero variance were excluded for LDA. Prediction analysis of microarrays was done with the pamr package *(57)*. Neural networks were built using Keras with a Tensorflow backend (10 layers, ~7×10^6^ parameters). Unless otherwise noted, default settings were used for tuning parameters as implemented in the respective packages.

### Differential expression analysis

For differential expression analysis of dataset 2 the R package limma was used *(58)*. A linear model was fit on the data with inclusion of the study as a factor. Differentially expressed genes were called using an FDR-corrected p-value < 0.05 and a minimum fold change of +/-2. For the permutation-based approach, 4174 samples were randomly drawn 100 times from the dataset. In each subset, DE genes were called as before, but without correcting for any batch in the model. The number of times each gene was called was summed up over all 100 permutations. Genes were ranked according to their overall DE count.

In addition to that a *l*_1_-penalized logistic regression was performed using the package glmnet *(53)* on the whole dataset and on each of the permutations. Genes were called to be of predictive importance if features had non-zero coefficients. The number of times each feature was of predictive importance was summed up, which resulted in a feature ranking of all “lasso genes”.

### Hierarchical Clustering

35 genes which had a stability of > 50% over 100 permutations were visualized using the R package pheatmap *(59)* (Fig. 4B). The data was z-scaled and columns clustered according to Euclidean distance. Rows were ordered according to diseases. Two gene clusters were visualized.

### Co-expression network analysis of transcriptional regulators

All 12.708 genes of dataset 2 were filtered for transcriptional regulators (TRs), resulting in 750 present TRs. Pearson correlation between genes was calculated using the rcorr function of the R package Hmisc *(60)*. 613 TRs with Pearson Correlation Coefficient > 0.509 and a p-value < 0.05 were visualized in a network with force-directed layout using Cytoscape. To define clusters within the network, all nodes were subjected to infomap community finding clustering *(61)* using the R package igraph *(62)*. Genes were clustered 1000 times and genes were allocated to the most frequent cluster. Genes that were advised to at least 20 different clusters were not assigned to any cluster and colored in white. This strategy resulted in 13 clusters, which were mapped on the network.

### Hierarchical clustering of co-expressed TRs

To find out whether clusters of co-expressed TRs were specific for distinct disease types, we plotted the mean fold change of all genes in one cluster vs. the corresponding overall mean in a heatmap. This was done for each disease individually and then visualized in one network graph using the R package pheatmap (56).

### Prediction of transcriptional regulators

To predict master regulators for genes of the AML-specific cluster, the 17 genes of the AML-specific cluster (*n*= 17 genes) were tested for enriched known transcription factor binding motifs using the function findMotifs.pl from the software package HOMER v4.9.1 *(63)* with standard settings and the promoter set “human”. The top ten best matches of enriched motifs were checked for presence in the co-expression network of TFs and visualized in the network.

## Acknowledgements

This work was supported by the German Research Foundation (J.L.S., SFB 704). J.L.S. is member of the Excellence Cluster ImmunoSensation. The research leading to these results received funding from the Horizon 2020 program (SYSCID Consortium). J.L.S is member of the Helmholtz network Sparse2Big.

